# Physical association of low density lipoprotein particles and extracellular vesicles unveiled by single particle analysis

**DOI:** 10.1101/2022.08.31.506022

**Authors:** Estefanía Lozano-Andrés, Agustin Enciso-Martinez, Abril Gijsbers, Sten F.W.M. Libregts, Cláudio Pinheiro, Guillaume Van Niel, An Hendrix, Peter J. Peters, Cees Otto, Ger J.A. Arkesteijn, Marca H.M. Wauben

## Abstract

Extracellular vesicles (EVs) in blood plasma are recognized as potential biomarkers for disease. Although blood plasma is easily obtainable, analysis of EVs at the single particle level is still challenging due to the biological complexity of this body fluid. Besides EVs, plasma contains different types of lipoproteins particles (LPPs), that outnumber EVs by orders of magnitude and which partially overlap in biophysical properties such as size, density and molecular makeup. Consequently, during EV isolation LPPs are often co-isolated. Furthermore, physical EV-LPP complexes have been observed in purified EV preparations. Since co-isolation or association of LPPs can impact single EV-based analysis and biomarker profiling, we investigated whether under physiological conditions LPPs and EVs can associate by using cryo-electron tomography, label-free synchronous Rayleigh and Raman scattering analysis of optically trapped particles and fluorescence-based high resolution single particle flow cytometric analysis. Furthermore, we evaluated the impact on flow cytometric analysis in the absence or presence of different types of LPPs using *in vitro* spike-in experiments of purified tumor cell line-derived EVs in different classes of purified human LPPs. Based on orthogonal single-particle analysis techniques we demonstrated that EV-LPP complexes can form under physiological conditions. Furthermore, we show that in fluorescence-based flow cytometric EV analysis staining of LPPs, as well as EV-LPP associations can influence EV analysis in a quantitative and qualitative manner. Our findings demonstrate that the biological colloidal matrix of the biofluid in which EVs reside impacts their buoyant density, size and/or refractive index (RI), which may have consequences for down-stream EV analysis.

## Introduction

Extracellular vesicles (EVs) are a heterogeneous group of membrane enclosed vesicles that contain biological information from the cell of origin, such as lipids, nucleic acids, carbohydrates and proteins, and are involved in intercellular communication. EVs present in blood plasma can be obtained via minimally invasive methods and have been proposed to hold clinical potential as biomarkers for diagnosis and prognosis of diseases since their specific makeup reflects a unique signature of the cell of origin [1-3]. Next to small EVs (50-200nm), plasma contains large amounts of lipoprotein particles (LPPs). LPPs are heterogeneous particles enclosed by a single layer of phospholipids and are often categorized based on their density and protein/lipid composition. The major LPP-types are chylomicrons (CM), very low-density lipoprotein (VLDL) particles, low-density lipoprotein (LDL) particles and high-density lipoprotein (HDL) particles [4]. CM have a size range of 75-1200 nm in diameter, with varying concentration between individuals and (fatty) meal consumption. VLDL are derived from CM, with 30-80 nm in diameter, which can be further transformed into LDL particles, with a smaller size diameter range of 5-35 nm. These LPPs have a low density (< 0.930-1.063 g/cm3) and are reported to contain copies of ApoB proteins [4, 5]. HDL particles are even smaller in diameter (size range 5-12 nm), have a high density (1.063-1.210 g/cm3) and do not contain ApoB proteins but ApoAI proteins [4, 5]. Recent studies have shown that the presence of LPPs is expected to interfere with both EV isolation and analysis, as they not only outnumber EVs by several orders of magnitude but also partially overlap in biophysical properties such as size and density [6-10]. The most widely applied EV isolation methods, differential (ultra)centrifugation, size-exclusion chromatography (SEC) and density gradient centrifugation have been reported to be unable to efficiently separate EVs from LPPs and described the co-isolation or formation of co-precipitates in the final preparations [7-9, 11-13]. For example, SEC is currently applied frequently for the analysis of clinical samples and although SEC allows separation of EVs and smaller HDL particles, EV-enriched SEC fractions still contain similarly sized ApoB+ particles [10]. This is in agreement with recent studies showing co-isolation between EVs and LPPs in fresh and processed blood plasma samples [8, 9, 14]. It has been demonstrated that the combination of different EV isolation methods can strongly reduce the amount of LPPs and can be applied to obtain EVs from blood plasma with high purity, however such procedures may also result in the selection of certain subpopulations of EVs [10, 15-18]. Moreover, the fact that the implementation of such combined isolation methods, is currently limited in a clinical setting, because these procedures are elaborate, time-consuming and expensive. Importantly, the fact that EV-LPP complexes have been demonstrated in EV preparations should be taken into account, since EV characteristics will be influenced in such complexes [8, 9]. Currently, it is not known whether EV-LPP complexes are artefacts resulting from the EV isolation methods used, or also occur in a physiological situation. Interestingly, some studies using minimal sample processing reported the presence of LPPs in close proximity to the lipid bilayer of EVs, indicating that EV-LPP interactions might indeed happen under physiological conditions [8, 9, 14]. Overall, this complex biological landscape complicates EV isolation and single EV-based analysis and limits the translation of EVs as biomarkers for diseases. Therefore, a better understanding on how the presence of distinct types of LPPs can affect the appearance of EVs is critical for the characterization of single EVs and their use as biomarkers in blood plasma. Techniques that allow for single particle analysis of heterogeneous populations are pivotal for this. Cryo-electron tomography (ET), although not being high-throughput, allows for the visualization of heterogeneous samples with great resolution and is able to distinguish between lipid bilayer EVs and single layer LPPs [19]. Label-free synchronous Rayleigh and Raman scattering analysis of optically trapped particles has been proposed as a feasible approach to detect and differentiate both EVs and LPPs based on their Raman spectrum and molecular composition with minimal need for sample processing [20]. Flow cytometry (FC) is a high-throughput multiparametric technique that is widely incorporated into clinical labs. However, detection of single EVs is challenging due to the resolution limit of most available instruments and the intrinsic features of EV, such as their small size and low refractive index (RI) [21, 22]. Previous studies have shown that the presence of LPPs in plasma can influence light scatter-triggered FC detection of EV [8, 23]. Fluorescence-triggered FC is an alternative for detection of EVs but depends on fluorescent staining procedures, e.g. staining with generic fluorescent dyes and/or incubation with fluorophore-conjugated antibodies against specific proteins [21, 22, 24-27]. We here used these three different analysis techniques to gain insight into the physiological interactions between LPPs and EVs. Furthermore, we evaluated the influence of LPPs on the quantitative and qualitative FC analysis of EVs and show the implications for EV-based biomarker profiling.

## Material and Methods

### Human lipoprotein particles

Purified human lipoprotein particles, i.e. human chylomicrons (0.89 mg/ml, catalog no. 7285-1000, Biovision Incorporated), human very-low-density and low-density lipoproteins (catalog no. 437647-5MG and 6 mg/ml, LP2-2MG, Merck Millipore, respectively) were purchased and stored according to the manufacturers’ instructions. For experimental condition we used a fixed volume of 2 μl from each sample.

### Human plasma

Blood samples from healthy human donors were collected in sodium citrate at a final concentration of 3.2% (0.105M). Platelet-depleted plasma (PDP) was obtained within 120 minutes after collection by two consecutive centrifugation steps at 2,500 x g for 15 minutes at room temperature. After each centrifugation step, the supernatant was transferred to a new sterile plastic tube and the pellet was discarded, after which depletion of platelets was verified with an hemato analyzer (0 × 10^4^ plt/µL). Samples were then transferred to 1.5 mL tubes and stored at -80°C until used. Collection of blood was approved by the Ethical Committee of Ghent University Hospital (approval EC/2014/0655). Participants provided written, informed consent.

### Preparation of EVs

4T1 murine mammary carcinoma cell line (American Type Culture Collection (ATCC), Manassas, VA) were used as a cellular source to obtain EVs. Cells were maintained in Dulbecco’s minimal essential medium (DMEM) supplemented with 10% fetal bovine serum, 100 U/mL penicillin and 100 μg/mL streptomycin (Invitrogen, Carlsbad, CA). Every month cell cultures were tested for Mycoplasma contamination using MycoAlert Plus kit (Lonza, Verviers, Belgium). EVs were prepared from conditioned medium (CM) of the 4T1 cell culture as previously described [16, 28, 29]. Briefly, cells were washed once with DMEM, followed by two washing steps with DMEM supplemented with 0.5% EV-depleted fetal bovine serum. Cells were then incubated at 37 °C and 5% CO2 with 15 mL DMEM containing 0.5% EV-depleted fetal bovine serum. After 24 h of culture and when cell confluency was > 70%, cell counting was performed with trypan blue staining to assess cell viability (>90%) using an automated cell counter (CountessTM, Thermo Fisher Scientific). Conditioned medium was then collected and centrifuged for 10 min at 300 *g* and 4 °C. The supernatant was passed through a 0.45 µm cellulose acetate filter (Corning, New York, USA) and concentrated at 4 °C approximately 300 times using a 10 kDa Centricon Plus-70 centrifugal unit (Merck Millipore, Billerica, Massachusetts, USA). After filtration through a 0.22 µm filter (Whatman, Dassel, Germany), concentrated conditioned medium was used for Optiprep density gradient ultracentrifugation. EVs were characterized following the MISEV criteria [30]. Optiprep (Axis-Shield, Oslo, Norway) density gradients (ODG) were prepared as previously described [28]. In brief, a discontinuous iodixanol gradient was prepared by layering 4 mL of 40%, 4 mL of 20%, 4 mL of 10%, and 3.5 mL of 5% iodixanol in a 16.8 mL open top polyallomer tube (Beckman Coulter, Fullerton, California, USA). One milliliter of concentrated conditioned medium was pipetted on top of the gradient and samples were centrifuged for 18 h at 100,000 ×*g* and 4 °C using a SW 32.1 Ti rotor (Beckman Coulter, Fullerton, California, USA). Fractions of 1 mL were collected from the top and EV-rich fractions 9 and 10 (corresponding to a density of 1.10-1.12 g/mL) were pooled for additional purification. Size-exclusion chromatography (SEC) was performed by using a nylon net with 20 µm pore size (NY2002500, Merck Millipore, Billerica, Massachusetts, USA) was placed on the bottom of a 10 mL syringe (BD Biosciences, San Jose, California, USA), followed by stacking of 10 mL Sepharose CL-2B (GE Healthcare, Uppsala, Sweden). On top of the SEC column, 2 mL of sample was loaded and eluted with PBS. Fractions of 1 mL were collected and EV-containing eluates 4-7 were pooled together. Pooled eluates were then concentrated approximately 40 times using a centrifugal filter (Amicon Ultra-2 10k, UFC201024, Merck Millipore, Billerica, Massachusetts, USA) following the manufacturers’ instructions. Concentrated EV-eluates were resuspended in PBS to a final volume of 100 µL and aliquoted in eppendorf tubes (20 µl each) and stored at -80 °C until further use.

### Dot blot analysis

Samples were spotted onto a 0.22 µm pore size nitrocellulose membrane (GE Healthcare) and allowed to dry. Membranes were subsequently blocked with PBS containing 0.5% (w/v) fish gelatin (Sigma-Aldrich) and 0.1% Tween-20 and incubated overnight at 4°C in a humidified chamber with primary human anti-CD9 (Biolegend, catalog no. 312102, dilution 1:1000), human anti-CD63 (BD, catalog no. 556019, dilution 1:1000) or human anti-ApoB100 (R&D Systems, catalog no. AF3260, dilution 1:1000). After washing with 0.1% Tween-20 in PBS, membranes were probed with secondary goat anti-mouse Polyclonal (Jackson ImmunoResearch catalog no.115-035-044, 1:10000) or donkey anti-goat Polyclonal antibodies conjugated with HPR (Invitrogen, catalog no. A16005, dilution 1:5000) and detected using Supersignal West Dura Extended Duration chemiluminscent substrate (Thermo Fisher Scientific). Imaging was performed using a ChemiDoc MP system and data was visualized using Image Lab Software v5.1 (Bio-Rad, Hercules, CA, USA).

### Electron Microscopy (TEM)

Purified human lipoprotein particles were deposited on carbonated grids and fixed in 2% PFA in 0.1 M phosphate buffer, pH 7.4. The grids were then embedded in methyl-cellulose/Uranyl Acetate 0,4%. All samples were examined with a FEI Tecnai Spirit electron microscope (FEI Company), and digital acquisitions were made with a numeric camera (Quemesa; Soft Imaging System).

### Synchronous Rayleigh and Raman scattering

For the optical setup measurements, and the characterization of single optically trapped particles, synchronous Rayleigh and Raman scattering acquisition was performed as described [20, 31] and briefly in Supporting Materials & Methods. The intensity and wavelength of the Rayleigh-Raman spectrometer were calibrated as described in [20] and briefly in Supporting Materials & Methods. Prior to the Rayleigh and Raman scattering measurements, all samples were diluted in PBS to prevent simultaneous trapping of multiple particles. Preparations of LPPs (CM, VLDL and LDLD), 4T1 EVs and EV-LPP mixtures were measured. For the LPP-EV mixtures, a fixed input volume of 5 μL of EVs, containing a nominal amount of 8.7E9 particles based on NTA analysis, were spiked with a fixed volume of 2 μL of LPPs (same as used from the samples analyzed in Figure 1a-c). The mixed samples were further diluted in PBS to a final volume of at least 300 µL and triplicates were measured alternating between sample types. For each sample type a volume of 50 µL was loaded in the well of a glass slide (BMS Microscopes; 1.0-1.2 mm thick), covered with a glass coverslip (VWR Ltd, thickness No. 1, diameter: 22 mm) and sealed with glue (EVO-STIK, Impact) to avoid evaporation. After the drying of the glue, each glass slide was placed under the microscope objective. The laser focal spot was focused inside the solution ⁓60 µm below the coverslip. In each cycle of 9.7 s, 256 Rayleigh-Raman spectra were acquired with an acquisition time of 38 ms per spectrum. The trapped particles were released from the laser focal spot by blocking the laser beam for 1 s. A total of 100 measurement cycles were acquired for each sample (n=13). Hence, a total of 332,800 Rayleigh-Raman spectra were acquired from which time traces were computed to identify the time intervals corresponding to individual trapping events. A total of 510 individual trapping events were analyzed (72 on average per sample type). The computation, segmentation and analysis of the Rayleigh and Raman time traces is described in detail in [20] and briefly in Supporting Materials & Methods.

**Figure 1.**
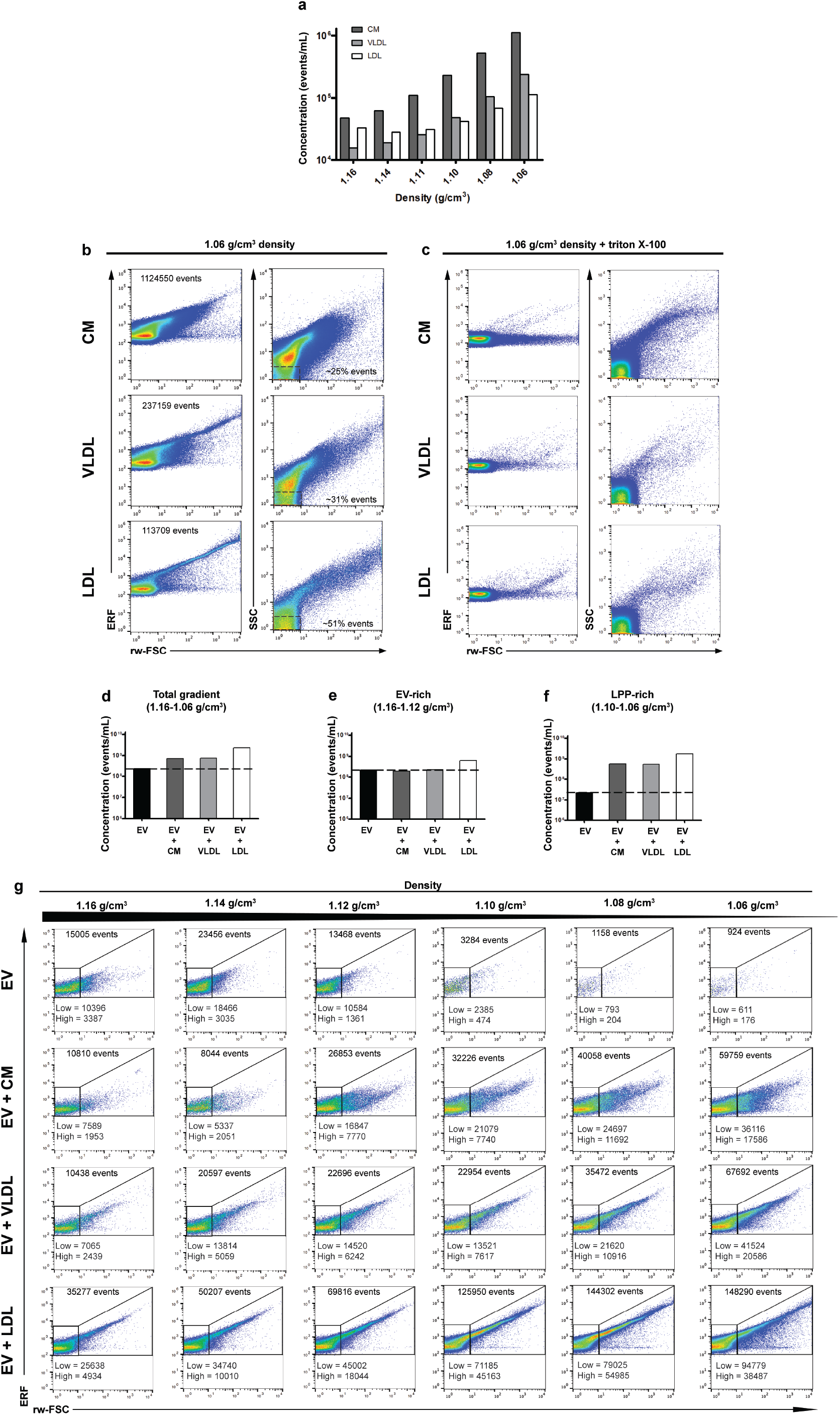
Analysis of generic fluorescence and light scattering profiles of commercial human LPP preparations and mouse EVs in presence or absence of LPPs. (a) Bar graph displaying the concentration of PKH67+ events in LPP, CM, VLDL and LDL preparations respectively, in each density fraction as determined using time-based flow cytometric analysis. (b) Representative dot plots displaying PKH67 fluorescence vs. reduced wide-angle FSC (rw-FSC) or SSC vs. rw-FSC of the low density fraction (1.06 g/cm3) from CM, VLDL and LDL preparations, respectively. Percentage of the gated low SSC vs. rw-FSC events from the total population is indicated. (c) Representative dot plots displaying PKH67 fluorescence vs. reduced wide-angle FSC (rw-FSC) or SSC vs. rw-FSC of the low density fraction (1.06 g/cm3) from LPPs after triton X-100 treatment (final concentration 0.1%). (d) Bar graph displaying total concentration of PKH67+ events in the density fractions of interest (1.06-1.16 g/cm3) (e) Bar graph displaying the total concentration of PKH67+ events in the EV-rich density fractions (1.12-1.16 g/cm3) or (f) in the LPP-rich density fractions (1.06-1.10 g/cm3) from the non-spiked, CM-, VLDL-or LDL-spiked EV samples, respectively. (g) Dot plots displaying PKH67 vs. rw-FSC of EV-rich density fractions (1.06-1.16 g/cm3) from the non-spiked, CM-, VLDL-or LDL-spiked EV samples, respectively. The total number of events for each plot is indicated on top of the plots. The number of the gated rw-FSC low or rw-FSC high populations is indicated within the plots.

### Cryo-electron tomography

The human lipoproteins were prepared in PBS/0.1% aggregate-depleted BSA, and BSA-gold 10 nm fiducials (OD600 1). A volume of 2.5 μL was applied on glow-discharged UltrAuFoil Au200 R2/2 grids (Quantifoil), and excess liquid was removed by blotting for 3 s (blot force 5) using filter paper followed by plunge freezing in liquid ethane using a FEI Vitrobot Mark IV at 100% humidity at 4 °C. Electron tomography data were acquired with a 200-kV Tecnai Arctica transmission electron microscope (Thermo Fisher Scientific) equipped with a Falcon III direct electron detector. Movies were acquired at 53k× magnification using a stage tilt scheme of −60° to 60° in increments of 3° through a total electron dose of 120 e −/Å2 and a defocus target range of −3 to −5 µm. Tilt series were aligned and reconstructed with IMOD using gold-particles tracking and SIRT, respectively [32].

### Fluorescent staining and labeling for high-resolution flow cytometric analysis

Generic staining of particles was performed as previously described [21] with some minor modifications indicated below. Briefly, 2 μL of CMs, VLDLs and LDLs or 5 μL of EVs were resuspended in 20 μL PBS/0.1% aggregate-depleted bovine serum album (BSA) prior to PKH67 staining (Sigma-Aldrich). The stock solution of aggregate-depleted BSA (5% w/v) was prepared by overnight centrifugation at 100,000 x *g* (SW28 rotor Beckman Coulter, Fullerton, California, USA; 4°C; κ-factor 334.2). For antibody labeling, samples were first resuspended in 20 μL PBS/0.1% aggregate-depleted BSA and incubated with 0.5 μg of Rat anti-mouse CD9-PE (Clone: KMC8, IgG2a, κ, Lot. no. 7268877, Becton Dickinson Biosciences) or matched Isotype antibodies (Rat IgG2a, κ, PE-conjugated, Lot. no. 8096525, Becton Dickinson Biosciences) for 1h at RT while protected from light exposure. After staining, samples were cleared from protein aggregates, unbound PKH67 dye and unbound antibodies by overnight bottom-up sucrose density gradient (SDG) ultracentrifugation at 192,000 *g* (SW40 rotor Beckman Coulter, Fullerton, California, USA; 4°C; κ-factor 144.5), according to the previously described protocol [21]. Gradient fractions of 1 mL were collected and densities were determined by using an Atago Illuminator (Japan) refractometer.

### High-resolution flow cytometric analysis

High-resolution flow cytometric analysis was performed with a jet-in-air-based flow cytometer (BD Influx, Beckton Dickinson Biosciences, San Jose (CA)) that is modified and optimized for detection of submicron-sized particles, and which is fully described in detail previously [21]. Upon acquisition, all scatter and fluorescence parameters were set to a logarithmic scale. To ensure that each measurement was comparable, a workspace with predefined gates and optimal PMT settings for the detection of 100 and 200 nm yellow-green (505/515) FluoSphere beads (Invitrogen, F8803 and F8848) was loaded. Upon aligning the fluid stream and lasers the 100 and 200 nm bead populations had to meet the criteria of pre-defined MFI and scatter values within these gates, where they displayed the smallest coefficient of variation (CV) for side scatter (SSC), reduced wide-angle forward scatter (rw-FSC) and FL-1 fluorescence. The trigger threshold level was set by running a clean PBS sample, thereby allowing an event rate ≤10-20 events/second. For detergent treatment of samples, 1% (v/v) triton X-100 (SERVA Electrophoresis GmbH, Heidelberg, Germany) was added to a final concentration of 0.1% triton X-100 and incubated for 30 seconds at RT prior re-analysis. When performing quantitative and qualitative analysis of submicron-sized particles, 50 μL of each fraction was diluted in 950 μL of PBS. Upon loading the sample, the sample was boosted into the flow cytometer until events appeared, after which the system was allowed to stabilize for 30 seconds. Measurements were then recorded for a fixed time of 30 seconds using BD FACS Sortware 1.01.654 (BD Biosciences). Particle concentrations were determined by measuring the flow rate of the instrument and correcting the number of detected particles for dilution and measured time. In between measurements of samples the sample line was washed subsequently with BD FACSRinse (BD Biosciences) and PBS for 5 seconds. Data analysis was performed using FlowJo Software version 10.0.8. Additional information according to the MIFlowCyt author checklist (Supplementary Table 1), MIFlowCyt-EV framework (Supplementary Table 2) and the calibration of the fluorescence axis (Supplementary Figure 1) using FlowJo Version 10.5.0 and FCMPASS Version v2.17 is provided in the Supplementary Material & Methods and described [33, 34].

## Data availability

We have submitted relevant data of our experiments to the EV-TRACK knowledgebase (ID: EV190078) [35]. Flow cytometry data files are available upon request.

## Results

### Fluorescence-triggered flow cytometry allows the detection of generic membrane-stained human LPPs in the absence or presence of EVs

To evaluate whether CMs, VLDLs and LDLs can be stained with generic membrane dyes and detected by fluorescence-triggered FC, we obtained commercially available purified human LPPs. To stain these LPPs, we used the lipophilic generic membrane dye PKH67, which has been successfully used for EV detection [21]. Based on its biochemical properties it is also expected that PKH67 can stain the single-layer enclosing the LPP particles. After incubation with PKH67 bottom-up sucrose density-gradient (SDG) ultracentrifugation was performed to evaluate the density at which the three types of LPPs could be detected. Fluorescence-triggered FC, showed for all LPP-types that the highest concentration of events was found in the lowest-density fraction (1.06 g/cm3) of the gradient (Figure 1a). The increase in detected events in the low-density fractions was as expected strongest for the CMs, which are the biggest and least dense LPPs and thus more easily pass the fluorescent threshold as compared to the smaller VLDL/LDL particles. Also at higher densities, corresponding to typical EV-densities (1.11-1.16 g/cm3), PKH67+ events could be detected, indicating that in the experimental set up used not all LPPs fully floated to their density equilibrium. Further analysis of the light scattering showed that the majority of CMs induced the strongest light scattering intensities (Figure 1b, upper panel), whereas LDLs, the smallest particles analyzed, induced the weakest light scattering intensities, with more than 50% of the particles displaying low light scattering signals (Figure 1b, bottom panel). Accordingly, VLDL particles, which fall in between CMs and LDLs in terms of size and heterogeneity, showed an intermediate light scattering profile (Figure 1b, middle panel). Although these observations are in agreement with the reported sizes of these LPPs, light scattering signals cannot be interpreted as a direct measurement of nanoparticle-size without appropriate calibration based on assumed RIs of these particles [36]. Interestingly, the LDL sample, and to a lesser extent the VLDL sample, showed a characteristic tail with increasing fluorescence, rw-FSC and SSC signals (Figure 1b, bottom and middle left panels), which can be generated by ‘swarming’ [37, 38] or by the presence of multi-particle LPP complexes.

Detergent lysis has been suggested as a control to confirm the detection of membrane enclosed EVs in samples and to rule out the detection of contaminants like protein complexes [39]. As LPPs display overlapping biochemical and biophysical features with EVs, we also evaluated the sensitivity of PKH67+ LPPs to detergent lysis. By incubating samples using triton X-100, a non-ionic detergent that has been reported to disrupt the lipid membrane of certain EV populations [39], we observed that upon re-analysis a great part of fluorescently stained LPPs, and/or their complexes disappeared upon triton X-100 lysis (Figure 1c). Indicating that these detergent lysis conditions cannot discriminate EVs and LPPs, which is consistent with previous reports also showing detergent lysis of LPPs under varying conditions (e.g. different staining and detection strategies) [6, 8]. After confirming that LPPs can be stained and detected with PKH67, we next investigated whether the presence of different types of LPPs affects the generic staining and detection of EVs. For this we purified EVs from the 4T1 murine mammary carcinoma cell culture supernatant by a combination of ODG ultracentrifugation and SEC (Supplementary Figure 1a). After EV isolation, EVs were characterized by Western blot, which confirmed the presence of transmembrane protein markers (e.g. tetraspanins CD9 and CD63), and by Nanoparticle Tracking analysis (NTA), which indicated a relative concentration of ∼1.8E12 particles/mL with a mean size distribution of ∼125 nm (Supplementary Material & Methods, Supplementary Figure 2b-c). For FC analysis EVs were stained with PKH67, followed by SDG ultracentrifugation. Time-based quantification of all gradient fractions showed that the peak density fraction of 1.14 g/cm3 contained the highest number of events for this 4T1EV preparation (Supplementary Figure 3a) and serial dilutions confirmed single EV detection (Supplementary Figure 3b). PKH67 fluorescence and light scattering signals from the EVs showed a clear and confined population (Supplementary Figure 3c).

After characterization of LPPs and EVs by FC, we next investigated how the simultaneous presence of LPPs and EVs in a preparation would affect the staining and/or detection. For this purpose we selected a fixed input volume of unstained 4T1 EVs (i.e., 5 ∼L containing a nominal amount of 8.7E9 particles based on NTA analysis) that was spiked with a fixed volume of LPPs (2 μL, similar as used in Figure 1a-c). Samples were next stained with PKH67 and fractionated by SDG ultracentrifugation for fluorescence-triggered FC analysis. The total number of PKH67+ events detected in all density fractions of interest, i.e. the sum of PKH67+ events in the range of 1.06 to 1.16 g/cm3 corresponding to both EV enriched fractions (1.12-1.16 g/cm3) and LPP-enriched fractions (1.06-1.10 g/cm3), was increased in the presence of LPPs (Figure 1d). Spiking CM or VLDL particles did not affect the total number of PKH67+ events in the EV-rich fractions (1.12-1.16 g/cm3), in contrast to spiking with LDL particles leading to an increase in the total number of PKH67+ events in these fractions (Figure 1e). Hence the presence of LDL particles affected fluorescence detection of EVs and thus quantitative EV analysis. In the LPP-rich fractions (1.06-1.10 g/cm3), the number of PKH67+ events increased in all spike-in samples (Figure 1f), whereas purified EVs displayed a very low number of events (Figure 1f), comparable to the procedural controls (Supplementary Figure 2c). Evaluation of the fluorescent and light scattering profiles of the EV-rich fractions revealed that LPPs also affect qualitative analysis by strongly increasing high light scattering signals in the spiked-in samples (Figure 1g). This was again most apparent in the presence of LDL particles. Overall, detected events in EV-LPP samples, in which multi-particle interactions might occur, exhibited higher fluorescent and light scattering intensity signals at different densities compared to EVs only. The PKH67 fluorescence and rw-FSC light scatter signals detected in the LPP-rich low-density fractions resembled the previously described PKH67+ LPP pattern (Figure 1b), while in the EV only sample very few events were detected in the low density fraction (Figure 1g, top row). Taken together, our findings demonstrate that the presence of CMs, VLDLs and LDLs in EV samples affects the quantitative and qualitative analysis of EVs.

### Association between human LPPs and EVs in physiological samples revealed by immunoblotting and cryo-electron tomography

The profiles from the LPPs revealed by FC, together with previous literature reports showing co-isolation of LPPs and EVs in EV preparations from blood samples [8-10], prompted us to also characterize the commercial LPP preparations in detail. As indicated by the manufacturer these LPP samples obtained from healthy human plasma donors have a > 95% purity, which we confirmed with transmission electron microscopy (TEM) showing spheroidal-shaped LPP particles (Figure 2). As expected the CM preparation contained relatively large particles with a rather heterogeneous size range (Figure 2a, upper row), LDL-samples contained the smallest particles with a fairly homogeneous size range distribution of approximately 20 nm (Figure 2a, bottom row), whereas VLDL particles were found to have a size range in between CM and LDL particles (Figure 2a, middle row). To evaluate whether these preparations contain a small portion of EVs, we next analysed the commercial LPP preparations also by immunoblot. We confirmed the presence of human Apolipoprotein B (ApoB), associated with these LPP types, in the stock solutions of the commercial LPPs and in a human platelet-depleted plasma (PDP) sample (Figure 2b). Since ApoB-48 is uniquely expressed in CM, whereas ApoB-100 is present in VLDL and LDL particles [4], the total ApoB signal cannot be used as an absolute measure for comparing these different samples. However, the volumetric-based analysis of the different preparations shows that the commercial LPP preparations contained less ApoB when compared to the physiological levels of ApoB detected in the PDP sample. Importantly, this indicates that the spike in effects of LPPs on FC analysis of EVs as observed in Figure 1, already occurred at relatively low LPP concentrations, and thus likely to happen as well in physiological blood plasma samples.

**Figure 2.**
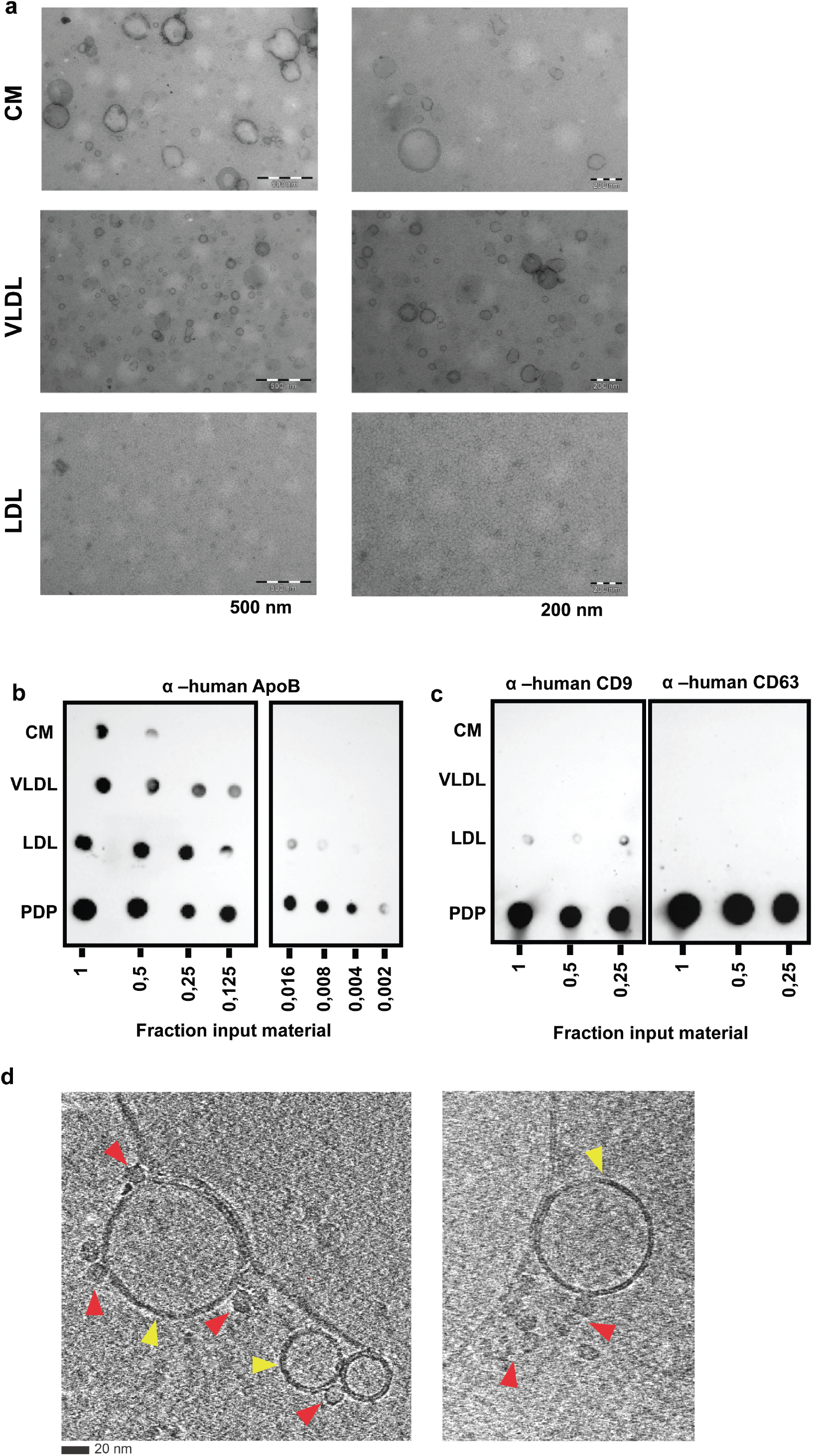
Analysis of purified LPPs by TEM, immunoblotting and cryo-electron tomography. (a) Analysis of human LPPs by TEM. 4 µl from a 1/10 dilution of commercial preparations of CM (top row), VLDL (middle row) and LDL (bottom row) particles were loaded onto grids, negatively stained and visualized. Scale bars correspond to 500 nm and 200 nm for each column. (b) Dot blot immunodetection of the commercial LPPs next to human platelet depleted plasma (PDP). Serial dilutions were spotted starting from 1 µl LPP stock or PDP and analyzed for the presence of ApoB by using a specific anti-human antibody against -ApoB. (c) Immunoblot detection of human tetraspanins - CD9 and -CD63 present in commercial LPPs preparations and in human PDP from a healthy donor.(d) Cryo-electron tomography of particles present in the commercial LDL preparation Lipid bilayer enclosed structures (i.e., EVs) are indicated with yellow arrows, while smaller lipid monolayer structures (i.e., LDLs) are indicated with red arrows. Bars = 20 nm for both images.

To detect the possible presence of EVs in the LPP preparations we used antibodies against two human tetraspanins present in plasma EVs (i.e., CD9 and CD63) for immunoblotting. As these tetraspanins are genuine transmembrane proteins, LPPs are negative for these proteins. However, a weak signal for human CD9 was detected in the commercial LDL preparation, but not in CM neither in VLDL preparations (Figure 2c). Based on the detection of this EV-marker in the LDL preparation we next used cryo-electron tomography to analyze possible EVs present in this preparation. With cryo-electron tomography single (i.e., LPPs) and double lipid layer (i.e., EVs) enclosed particles can be clearly distinguished. Besides the abundant present of LDL particles (∼ 20 nm) (Figure 2d, red arrows) we also identified the presence of bigger sized double lipid layered particles resembling EVs (Figure 2d, yellow arrows). Remarkably, these EVs were not randomly distributed along the grid, but often in close proximity to LDL particles forming complexes of EV-LDL. Furthermore, we also observed the presence of multi-LDL complexes that were forming clusters as their layers were physically in contact (Figure 2d, right picture).

### Label-free single particle synchronous Rayleigh and Raman scattering analysis unveiled the physiological formation of EV-LDL complexes

To investigate whether EV-LPP complexes can form in solution, we performed optical trapping of particles in suspension and acquired both Rayleigh and Raman scattering signals to detect individual trapping events and to characterize the particles chemical composition, respectively [20, 31]. We evaluated the scattering profiles of optically trapped particles present in the LPP (i.e., LDL, VLDL and CM) and 4T1 EV preparations, as well as in mixtures of LPP and EV preparations.

Individual trapping events were identified as a step-wise increase of the Rayleigh signal when plotted over time. By segmenting individual trapping events a Raman spectrum per trapping event was obtained, which was corrected by background subtraction [20, 31]. To compare Raman spectra of trapped particles, principal component analysis (PCA) was performed on the 4T1 EV preparation, the different LPP preparations, and the EV-LPP mixtures (Figure 3a-c). Whereas, the particles trapped in the EV-CM mixture (Figure 3a) or in the EV-VLDL mixture (Figure 3b) clustered together with respectively CM or VLDL particles only, particles present in preparations of LDL, 4T1 EVs and a mixture of LDL and 4T1 EVs clustered in three separate groups (Figure 3c). Since particles clustering together have similar chemical composition, this indicates that we mainly trapped VLDL or CM particles in the EV-VLDL mixture and the EV-CM mixture, respectively. In contrast, particles in the EV-LDL mixture neither overlap with particles trapped in the LDL preparation, nor with particles trapped in the EV preparation, suggesting that the particles trapped in the mixed sample have a different chemical composition As shown in Figure 3c, particles trapped in the EV-LDL mixture have PC1 scores >0, similar to LDL particles, and also mainly PC2 scores >0, resembling the EV sample. This suggests that Raman features from both LDL and EVs contribute to the Raman spectrum of the particles trapped in the mixed sample. Since the difference between LDL and EVs was highest for PC1, we next analyzed the PC1 loading to identify the source of these differences. Figure 3d shows PC1 loading displaying positive and negative peaks at wavenumber positions that correspond to triglycerides (green lines) and cholesterol (red lines), respectively. This means that particles with positive PC1 scores, such as particles trapped in the LDL preparation and in the LDL-EV mixture, have higher triglyceride and less cholesterol contributions to their Raman spectrum than particles with negative PC1 scores, such as particles trapped in the EV only preparation. Detailed analysis of the mean Raman spectrum between 1600 and 1700 cm-1, shows that particles trapped in the EV preparation contain only cholesterol (1670 cm-1), while particles present in the LDL preparation predominantly contain triglycerides (1657 cm-1) (Figure 3e). The presence of cholesterol in the EV preparation was further confirmed by Raman bands associated to cholesterol in the high frequency region (2850, 2866, 2888 and 2932 cm-1) (Supplementary Figure 4). Interestingly, particles trapped in the EV-LDL mixture showed clear contributions of both triglycerides and cholesterol in the Raman profile (Figure 3e). Importantly, these results show that EV-LPP associations are not only induced by isolation and/or staining procedures, but also form spontaneously in solution with label-free particles. Interestingly, LDL particles are more prone to form complexes with EVs than VLDL and CM.

**Figure 3.**
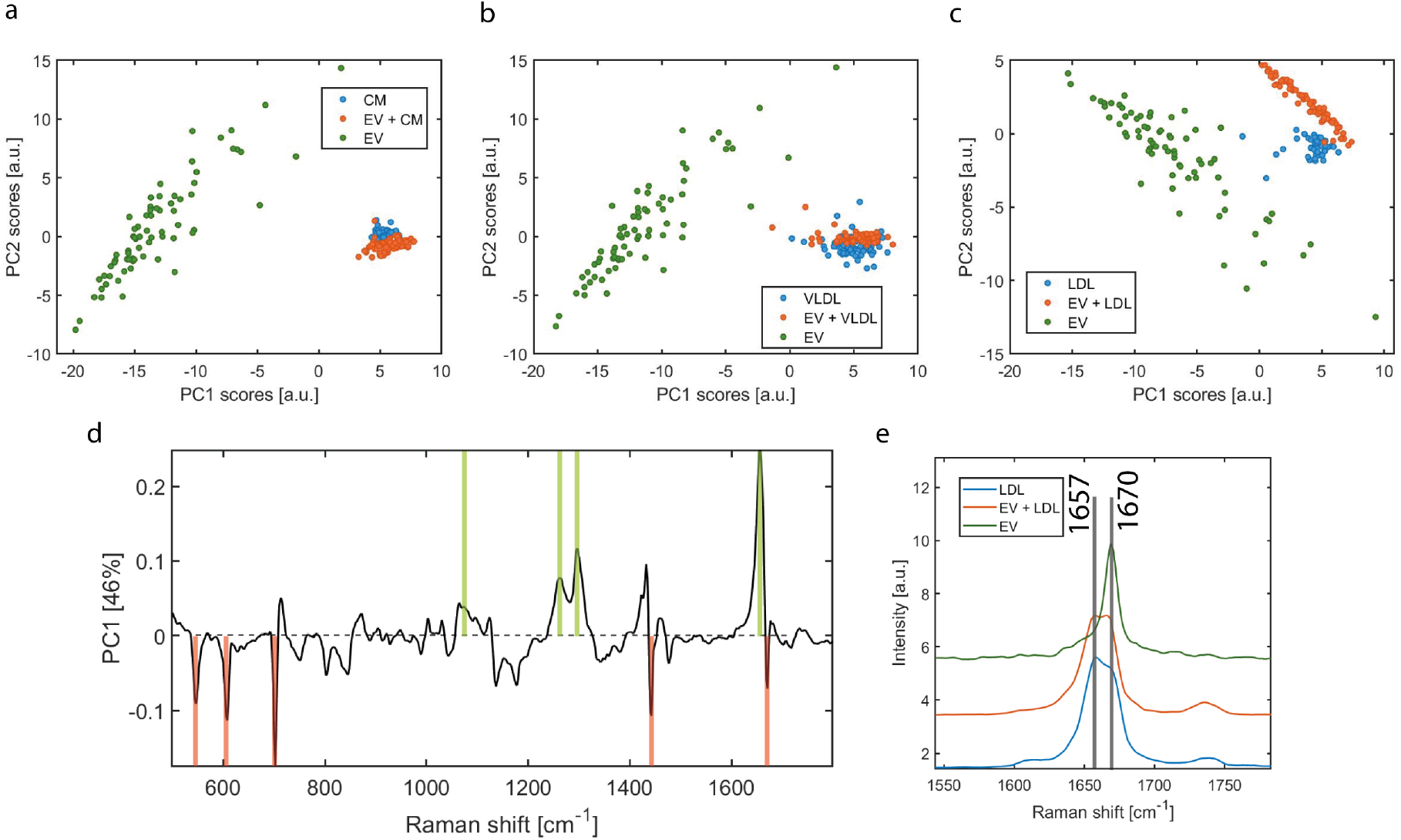
Principal component analysis (PCA) of individual LPP and EV preparations and mixed LPP and EV preparations. First and second principal component scores corresponding to the PCA of (a) CM, EV + CM and EV samples, (b) VLDL, EV + VLDL and EV, and (c) LDL, EV + LDL and EV (d) First principal component loading corresponding to the PCA of LDL, EV + LDL and EV. The green lines indicate triglyceride bands and the red lines indicate cholesterol bands. Particles with positive PC1 scores values in (c) are positively correlated with triglycerides, while negatively correlated with cholesterol. (e) Mean Raman spectrum per sample type (LDL, EV + LDL, EV) displaying a part of the Raman fingerprint region that shows triglyceride (1657) and cholesterol (1670) Raman bands.

### EV-LPP associations can impact the identification and detection of specific EV markers

Since we found that EV-LPP associations can form spontaneously in solution, implicating that EV-LPP complexes are likely present in blood plasma, we further investigated whether such associations can impact the identification and detection of specific EV-markers by FC. Our experimental set-up in which mouse-derived 4T1 EVs were spiked-in human LPP preparations allowed the use of a murine-specific CD9 antibody to detect the EV-marker of interest. We confirmed the exclusive detection of murine CD9+ EVs without cross-reactivity in human LPP preparations by immunoblotting (Supplementary Figure 2d), thereby excluding the risk of the detection of human CD9+ EVs already present in the human LPP preparation, as shown before (Figure 2c). Specific FC detection of purified murine PKH67+CD9+ EVs was confirmed (Supplementary Figure 3), and the fluorescent intensity was calibrated (Supplementary Figure 1).

When equal numbers of murine EVs were stained for PKH67 and murine CD9 in the absence or presence of LPPs followed by density gradient centrifugation, different numbers of CD9+ events were detected in time-based quantitative EV measurements of the total density gradient fractions (1.06-1.16 g/cm3) (Figure 4a). In the presence of LDL particles, the total number of CD9+ events was strongly increased, while no or marginal increase of the number of CD9+ events was observed in the presence of CM or VLDL particles, respectively (Figure 4a). In contrast, in the EV-rich density gradient fractions (1.12-1.16 g/cm3) the highest number of CD9+ EVs was detected in the EV sample in the absence of LPPs and the presence of CM and VLDL substantially reduced the number of CD9+ events(Figure 4b). Previously, we showed that the number of PKH67+ events in EV-rich densities were consistent for EVs in the absence or presence of CM or VLDL(Figure 1e). Hence, the detection of 2-fold less CD9+ EV in the presence of CM and VLDL (Figure 4b and 4d) indicates that CM and VLDL significantly affected the labelling and/or detection of CD9+ EVs. Although in the presence of LDL the number of CD9+ events was only marginally decreased (Figure 4b), the % reduction of CD9+ events was strongest (Figure 4d). This reduction in % CD9+ events is caused by the fact that in the presence of LDL the total number of PKH67+ events was increased in EV-rich fractions (Figure 1e), in contrast to the number of PKH67+ events of EVs in the presence of CM or VLDL that was similar to EVs only (Figure 1e).

**Figure 4.**
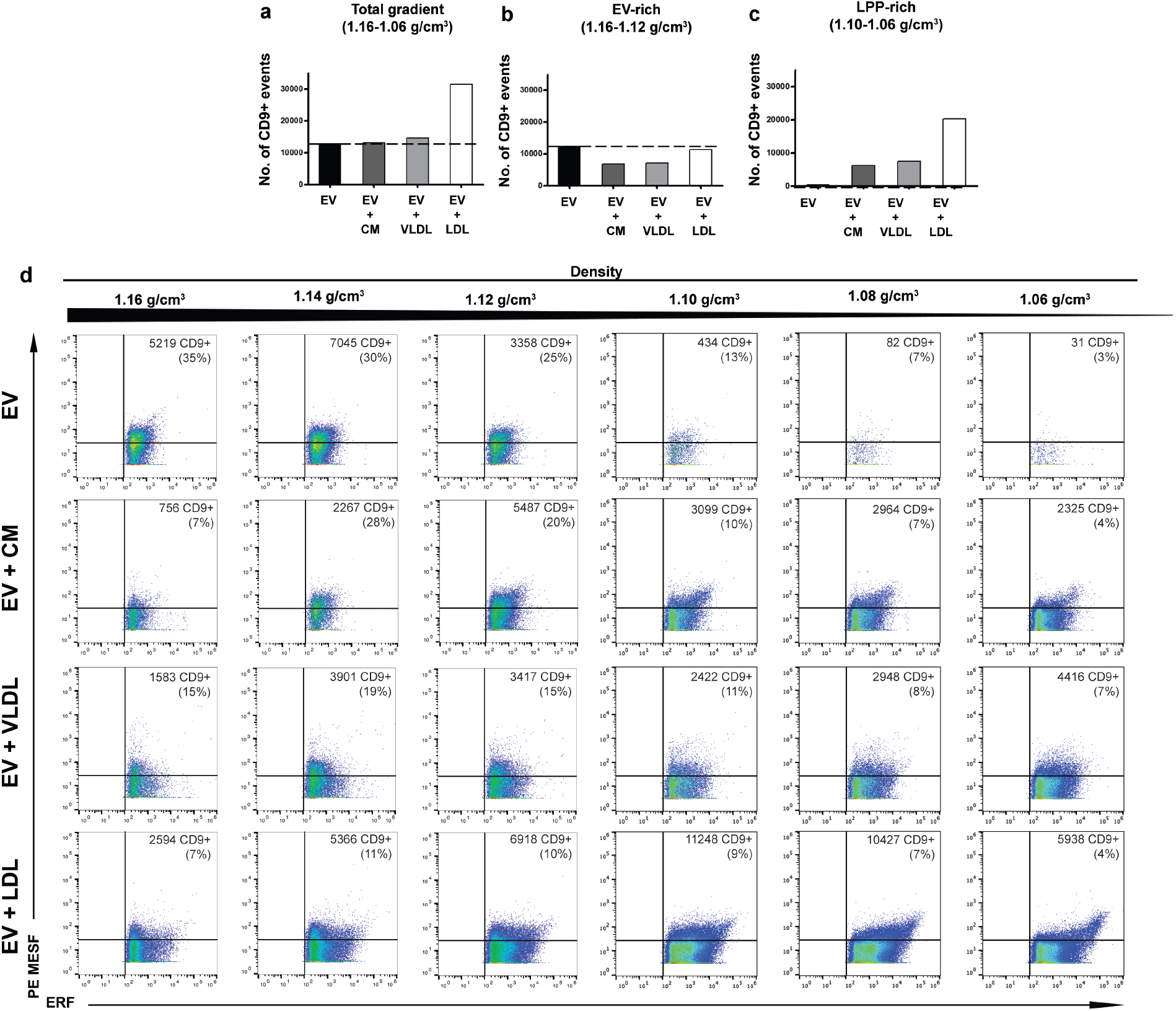
Detection of murine CD9+ EVs in the absence or presence of human CM, VLDL or LDL particles by fluorescence-triggered high-resolution flow cytometry. (a) Bar graphs displaying the number of PKH67+ CD9+ events in all density fractions of interest (1.16-1.06 g/cm3), (b) Bar graphs displaying the number of PKH67+CD9+ events in EV-rich density fractions (1.16-1.12 g/cm3) or (c) in the LPP-rich density fractions (1.10-1.06 g/cm3) from murine EVs in the absence or presence of CM, VLDL or LDL particles. (d) Dot plots displaying CD9+PE events in PE MESF units vs. PKH67+ events in ERF units from murine EVs in the absence or presence of CM, VLDL or LDL particles. The number of gated CD9+ events and its percentage from the total population is indicated for each plot.

Strikingly, a strong increase in the number of CD9+ events was observed in the LPP-rich densities (1.06-1.10 g/cm3) when EVs were analyzed in the presence of LLPs, with the strongest effect in the presence of LDL (Figure 4c). Analysis of the corresponding dot plots revealed that CD9+ events, especially in the presence of LDL had a distinct profile (i.e., displaying higher PKH67+ and CD9+ intensities) (Figure 4d), which could be indicative for physical EV-LDL interactions. Altogether, these data demonstrate that the co-presence of LPPs in EV-samples, not only affects generic EV staining but can also impact the staining, enumeration and buoyant density of EV-subsets labeled for specific EV-markers.

## Discussion

To exploit the use of EV-based biomarkers in blood confounding effects of non-EV components, such as the abundant and variable presence of LPPs, need to be considered. Circulating EVs are often analyzed by FC and recent studies indicated confounding effects of LPPs present in EV preparations in flow cytometric EV analysis [6-8]. Based on the partially overlapping properties of EVs and LPPs, optimized sequential biophysical fractionation protocols were developed to separate LPPs from EVs [10, 15, 17]. However, in recent studies it has been indicated that also complexes of EV-LPP can be detected in EV samples [8, 9, 40].

We here explored whether such EV-LPP complexes can be formed in solution, or are merely a result of the EV isolation and/or labeling methods used, and how LPPs and EV-LPP complexes affect downstream EV analysis by FC. By using the generic membrane dye PKH67, known to be incorporated by a wide variety of lipid-containing components due to its lipophilic nature [41], we stained highly purified LPPs and EVs which resulted in partially overlapping fluorescent and light scattering signals as measured by FC.

However the majority of fluorescent events was detected at different buoyant densities (i.e, LPP-rich density fractions (1.06-1.10 g/cm3 and EV-rich density fractions (1.12-1.16 g/cm3). Furthermore, in agreement with previous observations our data show that treatment with triton X-100 could not distinguish between LPPs and EVs, as both showed sensitivity to lysis under the exact same treatment conditions [8]. Previously, we optimized the PKH67 staining protocol for single EV-based FC and demonstrated the need of density gradient centrifugation to get rid of PKH-aggregates, which reside in high density gradient fractions excluded from our FC analysis [21]. Moreover, procedural controls were included to evaluate possible contributions of PKH67 artifacts. Using a fluorescence-based threshold triggering FC approach, the detection limit of particles depends on their generic fluorescent staining (PKH67) intensity, allowing only the detection of particles that exceed the set threshold, which in our instrument was calibrated to ~100 ERF units based on FITC MESF beads [42, 43]. Due to their size and refractive index, single VLDL and LDL particles are unlikely to be fully resolved at the single particle level with the settings that we used for FC analysis. However, their aggregates or multi-particle complexes can be detected by FC [8], and various reports have investigated and confirmed the formation of such physical LPP aggregates both *in vitro* and *in vivo* [44, 45]. Overall, our results clearly indicate the need for specific markers to attribute PKH67+ fluorescent events to LPPs or EVs, which is in accordance to previous observations [41, 46].

Our spike-in experiments using purified human LPPs and murine EVs were designed as a proof-of-principle study to address the knowledge gap on the impact of LPPs on EV analysis and to specifically investigate to formation of EV-LPP complexes [7]. To avoid artefacts resulting from too high concentrations of LPPs in these spike-in experiments, we intentionally used lower amounts of purified LPPs as compared to platelet depleted plasma samples (determined by immunoblotting for ApoB). Since we did not found a reduction of the number of PKH67+ events in the EV-rich fractions, while a strong increase in PKH67+ events was observed in the LPP-rich fractions after EV-LPP spike-in, we do not have indications that the PKH dye was a limiting factor resulting in reduced staining efficiency of EVs in the EV-LPP sample. Importantly, the flow cytometric analysis of a specific EV marker (in our experimental set-up murine CD9) on EVs in the presence of LPPs clearly demonstrated the presence of murine EV-human LPP complexes in these samples. Consistent with our observations, also others have found EV-associated markers at lower densities as expected in blood plasma [10]. Furthermore, we detected in the purified human LDL preparation weak human CD9 signals by immunoblotting, and confirmed by cryo-ET the occurrence of EVs in this LDL preparation, decorated or surrounded by multiple LDL particles. Also others previously demonstrated the presence of CD9 in purified LPP preparations (i.e., purified HDL) [7]. Importantly, our findings are in line with recent reports showing that interactions between EVs and ApoB-containing LPPs can be observed using various isolation methods and detection techniques [11, 13]. However, since these experiments cannot rule out that the formation of LPP-EV complexes might be induced by the particle isolation, purification and/or staining methods used, we also explored the formation of such complexes in solution. We here demonstrate for the first time, by using label-free single particle synchronous Rayleigh and Raman scattering analysis of purified LPPs, purified EVs and their mixtures, that complexes between EVs and especially LDL particles form spontaneous in solution.

Follow-up studies are needed to address the amount of EV-LPP complexes in the blood circulation and to define EV and LPP subsets prone to form such complexes in health and disease conditions. For EV-based biomarker profiling in blood samples, our current findings demonstrate the importance of critical sample collection, preparation and experimental design, since the presence of EV-LPP complexes can impact the detection of EV markers, which can hamper but can also be exploited to identify EV-subsets of interest.

## Funding Sources and acknowledgements

E.L.A. and C.P. are supported by the European Union’s Horizon 2020 research and innovation programme under the Marie Skłodowska-Curie grant agreement No 722148 (TRAIN-EV). S.F.W.M.L. was supported by the Dutch Technology Foundation STW (Perspectief Program Cancer ID, project 14191), which is part of the Netherlands Organisation for Scientific Research (NWO), and which is partly funded by the Ministry of Economic Affairs. G.v.N. is supported by Fondation pour la Recherche Médicale (AJE20160635884) Institut National du Cancer (INCA N°2019-1033125 PLBIO19-059). We thank Laura Varela-Pinzon (Department of Biomolecular Health Sciences and Department of Clinical Sciences, Faculty of Veterinary Medicine, Utrecht University) for valuable discussions and lipidomic analysis of LPPs. E.L.A. and M.H.M.W greatly acknowledge the FACS Facility at the Faculty of Veterinary Medicine at Utrecht University and G.v.N. greatly acknowledge the Electron microscopy facility of the Institut Curie.

## Author contributions

E.L.A. designed and performed experiments, analyzed data and wrote the original manuscript. A.E.M, A.G, S.F.W.M.L., C.P., and G.v.N. performed experiments, analyzed data and reviewed the manuscript. A.H., P.J.P and C.O. gave conceptual advice and reviewed the manuscript. G.J.A.A. supervised and designed the flow cytometric experiments and wrote the original manuscript. M.H.M.W. supervised the research, designed experiments and wrote the original manuscript. All authors critically reviewed and edited the manuscript.

