## Supporting Information for "Physical association of low density lipoprotein particles and extracellular vesicles unveiled by single particle analysis"

### Supplementary Material & Methods

**Western blot analysis of EVs.** Isolated EVs (1 µL or 4 µL) were resuspended in 3 µL PBS and 11 µL non-reducing SDS-PAGE sample buffer or 11 µL of non-reducing SDS-PAGE sample buffer, respectively. Samples were next separated on a 12.5% SDS-PAGE gel and transferred to a PVDF-membrane with a 0.45 µm pore size (Millipore, Amsterdam-Zuidoost, the Netherlands). Membranes were blocked for 1 hour using PBS containing 0.5% (w/v) fish gelatin (Sigma-Aldrich) and 0.1% Tween-20 and subsequently immunolabeled overnight at 4°C with primary antibody Rat anti-mouse CD9 (eBioscience, Thermo Fisher Scientific, Waltham, MA, USA; Clone KMC8, Lot #E028516, catalog #14-0091-85, 0.5 mg/ml, dilution 1:1000) or Rat anti-mouse CD63 (Biolegend, San Diego, CA, USA; Clone NVG-2, catalog #143902, dilution 1:500). After washing, membranes were incubated with a secondary antibody rabbit anti-rat PolyClonal conjugated to horseradish peroxidase (Jackson ImmunoResearch, Ely, Cambridgeshire, UK; 1:5000). Peroxidase was detected using Supersignal West Dura Extended Duration chemiluminscent substrate (Thermo Fisher Scientific). Imaging was performed using a

ChemiDoc MP system and data was visualized using Image Lab Software v5.1 (Bio-Rad, Hercules, CA, USA).

**Nanoparticle tracking analysis (NTA) of EVs.** Size distribution and concentration of isolated EV were determined by measuring the rate of Brownian motion using a NanoSight LM10 system microscope (NanoSight Ltd, Amesbury, UK) equipped with a camera sCMOS and a 405 nm laser. An automatic pumping system was used for the injection of the sample. For each sample, 3 videos of 30 seconds were recorded and analysed with a detection threshold 3 and camera level 13. The measurements were performed and monitored at ambient temperature which did not exceed 25°C. Recorded videos were then analysed with NTA Software version 3.2. For optimal measurements, samples were diluted in PBS until particle concentration was within the dynamic range of NTA Software (between 3E8 and 5E9 particles/ml).

**Fluorescence calibration.** For calibration of the fluorescence axis, FITC MESF beads (custom made, lot MM2307#131-10; #131-8; #130-6; #130-5; 130-3, BD Biosciences, San Jose, CA) and PE MESF beads (Lot. No. 62981, BD Biosciences, San Jose, CA) were used according to the manufacturer's instructions. Calibration MESF beads were provided with assigned numbers of molecules of equivalent soluble fluorochrome (MESF) for each peak intensity. Briefly, each bead peak population was gated using FlowJo Version 10.5.0 and median fluorescence intensities (MFI) were obtained (Supplementary Figure 4). Next, FCMPASS Version v2.17 was used to perform a least-square linear regression analysis and generate the data displaying MESF calibrated axis.

**Synchronous Rayleigh and Raman scattering.** A Rayleigh-Raman spectrometer was used, which is based on a home-built Raman spectrometer integrated with the base of an upright optical microscope (Olympus BX41). For illuminating and trapping the particles, a single laser beam from a Coherent Innova 70C laser ($\lambda_{exc}$=647.089 nm) was used. A cover glass corrected dry objective (Olympus, 40x, NA: 0.95) was used to collect the Rayleigh and Raman scattering, which was then separated in a homebuilt spectrometer and simultaneously detected with a single CCD camera (Andor Newton DU-970-BV). The average dispersion of the spectrometer was ~2.3 $cm^{-1}$ (0.11 nm) wavenumber per pixel over the CCD camera surface with 1600 pixels along the dispersive axis and 200 pixels along the other axis. The laser power was measured underneath the objective and adjusted to 70 mW.

**Calibration of the Rayleigh-Raman spectrometer.** Both intensity and wavelength calibration were performed to convert the raw measured data from pixels vs relative counts to calibrated data in wavenumber ($cm^{-1}$) vs counts. The pixel-to-wave number conversion was performed by using the toluene spectrum and an argon-mercury lamp with narrow emission lines known with picometer accuracy. The intensity calibration was performed by acquiring a white light spectrum from a tungsten halogen light source (AvaLight-Hal; Avantes BV, Apeldoorn, The Netherlands) that resulted in the light detection efficiency per pixel for the entire setup. The offset of the detector was obtained from a measurement without any light falling onto the detector. To determine the background spectrum, the spectrum of the entire light path through the setup with the laser on, but without a sample, was acquired and subtracted from the measured data. All software was developed in-house with Matlab (2017b, MathWorks, USA).

**Rayleigh-Raman data analysis.** Rayleigh and Raman time traces were obtained by integrating the Rayleigh (-20 – 10 $cm^{-1}$) and lipid-protein Raman bands (2811 – 3023 $cm^{-1}$), respectively. The integrated Rayleigh and Raman values were concatenated and plotted against time to enable the selection of individual trapping events visualized as step-wise increases in the Rayleigh signal [1, 2]. These steps were manually segmented and their time intervals were used to obtain their corresponding Raman spectra. A Raman spectrum of a single particle was thereby acquired as the average Raman spectrum along the length of each segmented step. To remove the background contribution, e.g. water or soluble components, also present in the focal spot with each single trapped particle, the immediate previous segment to each step was segmented and its corresponding Raman spectrum was used for background subtraction using linear least squares fit. The mean Raman spectrum of each particle was interpolated to the same wavenumber axis and baseline correction and de-noising were performed on the full dataset using "Baseline Estimation And Denoising with Sparsity" [3]. Each Raman spectrum was then normalized to have a mean of zero and a standard deviation of one. Principal component analysis was performed in the Raman fingerprint spectral region (500 – 1800 $cm^{-1}$). All software was developed in-house in Matlab (2017, MathWorks, USA).

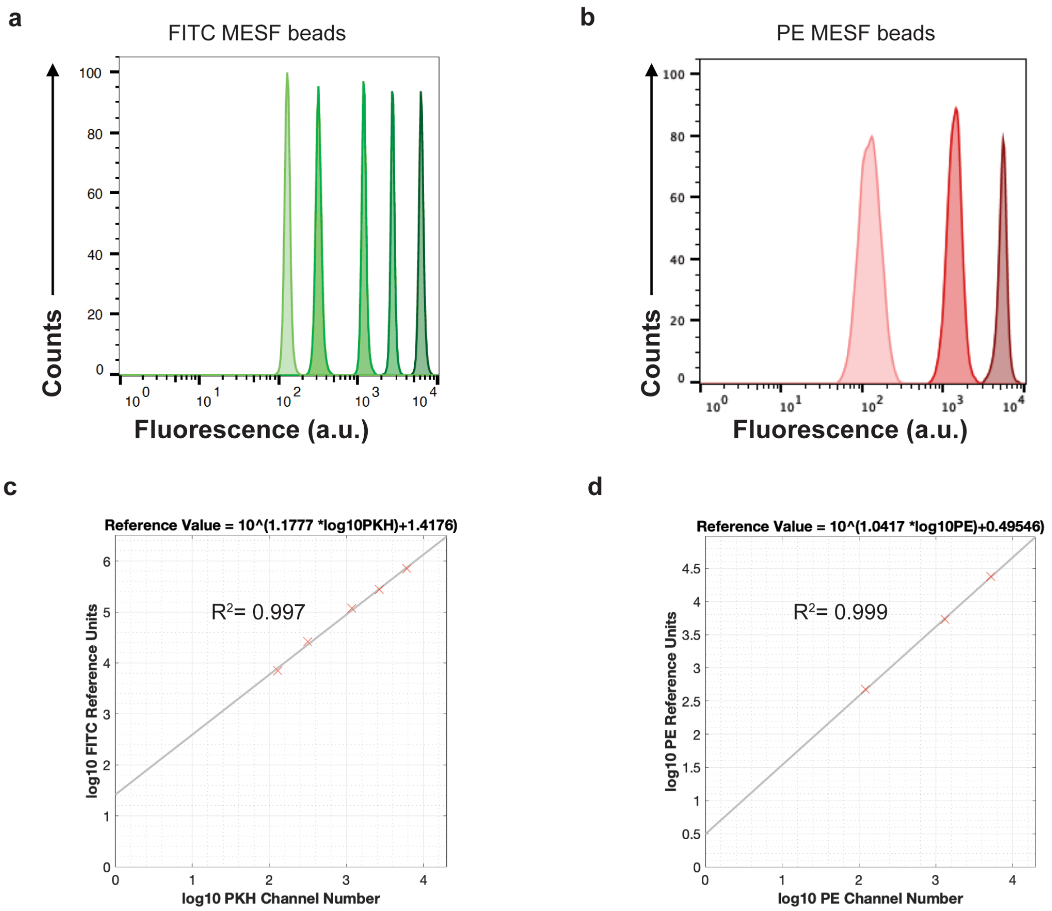


**e**

| Peak | MFI | LogMFI | FITC MESF | LogFITC MESF |
|---|---|---|---|---|
| P1 | 129 | 2.1106 | 7094 | 3.8508 |
| P2 | 317 | 2.5011 | 25910 | 4.4134 |
| P3 | 1173 | 3.0693 | 118192 | 5.0725 |
| P4 | 2684 | 3.4288 | 275288 | 5.4397 |
| P5 | 6111 | 3.7861 | 715225 | 5.8544 |

**f**

| Peak | MFI | LogMFI | PE MESF | LogPE MESF |
|---|---|---|---|---|
| P1 | 122 | 2.0863 | 474 | 2.6757 |
| P2 | 1325 | 3.1222 | 5359 | 3.7290 |
| P3 | 5186 | 3.7148 | 23843 | 4.3773 |

**Supplementary Figure 1.** Standardization of fluorescence intensities by using FITC and PE MESF calibration beads. (a) Histogram overlay of FITC-MESF beads and (b) PE-MESF beads. Axis is in arbitrary fluorescence units. (c) Relation between log10 fluorescence channel numbers and log10 FITC-MESF values or (d) PE-MESF values. (e) Characterization of the FITC-MESF beads and (f) PE-MESF beads. Shown are the respective peak numbers with their corresponding MFI as measured on the BD Influx, the log10 conversion thereof, the number of FITC-MESF or PE-MESF per bead and the log10 conversion thereof.

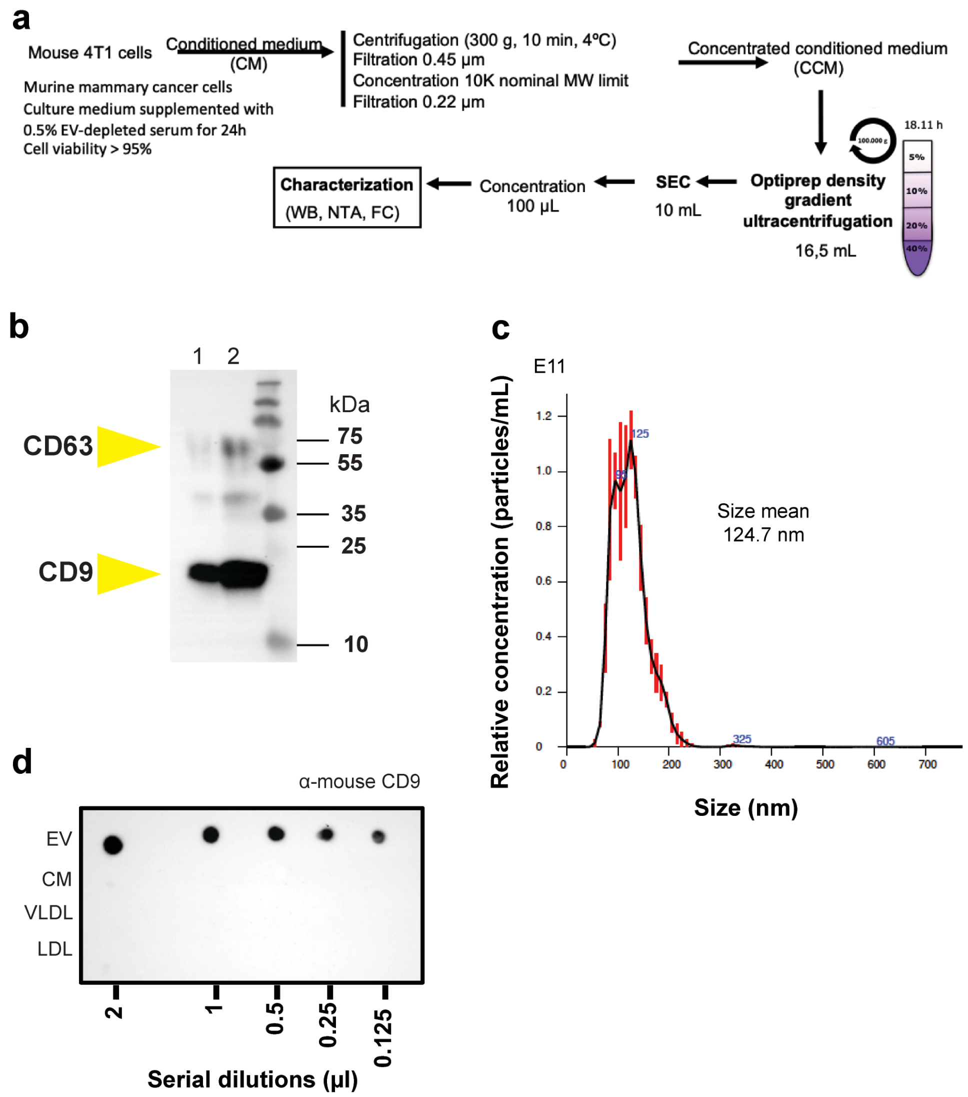


**Supplementary Figure 2.** Isolation, purification and characterization of mouse 4T1 EVs. (a) Schematic representation of mouse 4T1 EV isolation and purification methods. (b) Western blot analysis of purified EV. Samples were subjected to SDS-PAGE and detected by chemiluminescence using specific mAbs directed against mouse -CD9 (clone KMC8) and -CD63 (clone NVG-2). Lane 1 is a ¼ dilution from input in lane 2. (c) Nanoparticle tracking analysis (NTA) of purified EVs for size determination and quantification. (d)

Side-by-side dot blot detection of mouse CD9 in mouse 4T1 EVs and commercial LPP samples.

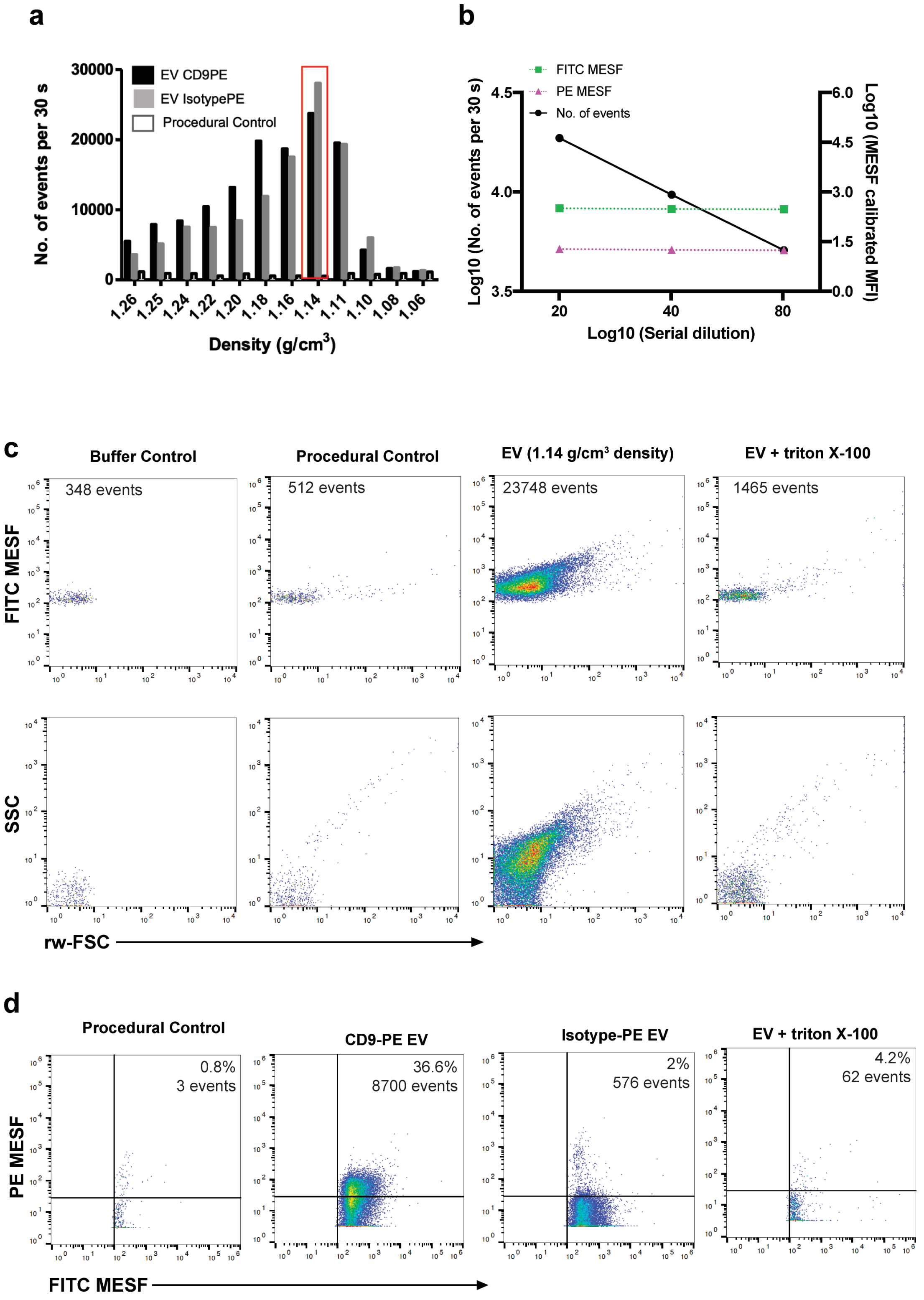

**Supplementary Figure 3.** Immunocharacterisation of mouse 4T1 EVs by high-resolution flow cytometry. (a) Bar graphs displaying the fractionation profile of stained EVs after sucrose density gradient floatation. The number of events in each density fraction is determined using time-based flow cytometric analysis (number of PKH67-positive events in 30 seconds). (b) Serial dilutions (1:20, 1:40, 1:80) of the fraction containing CD9 and PKH67 stained EV were analysed. Total number of events and MFI for FITC MESF and PE MESF are shown. (c) Dot plots of FITC MESF vs rw-FSC, SSC vs rw-FSC for experimental controls and the peak EV-fraction showing sensitivity of samples to detergent treatment (Triton X-100; final concentration of 0.1%). Respective total number of events are shown in each plot. (d) Dot plots of PE MESF vs FITC MESF for relevant experimental controls and the peak EV-fraction.

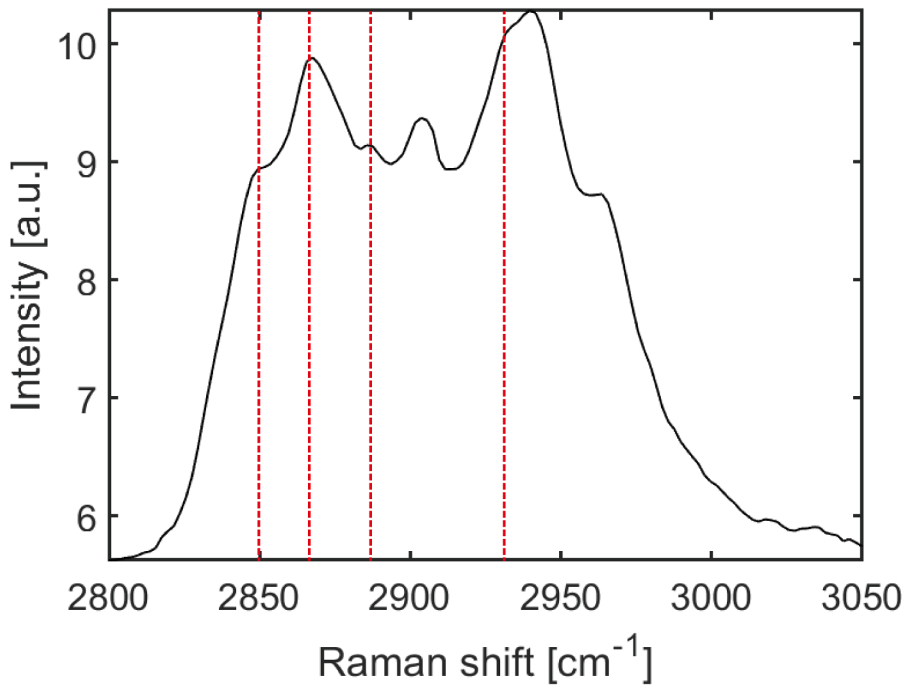

**Supplementary Figure 4.** High frequency region of the mean Raman spectrum of 4T1 EV. Red lines indicate Raman bands associated to cholesterol (2850, 2866, 2888 and 2932 cm-1).

**Supplementary Table 1. Author Checklist: MIFlowCyt-Compliant Items.**

| **Requirement** | **Please Include Requested Information** |
|---|---|
| 1.1. Purpose | Evaluate the influence of lipoprotein particles (LPP) for Extracellular Vesicle (EV) detection upon fluorescent threshold triggering by using the generic EV marker PKH67 upon fluorescence-based flow cytometry detection. |
| 1.2. Keywords | extracellular vesicles, lipoprotein particles, flow cytometry, plasma, biomarker |
| 1.3. Experiment variables | Different types of lipoprotein particles (Chylomicrons, VLDL and LDL) |
| 1.4. Organization name and address | Department of Biochemistry & Cell Biology<br>Faculty of Veterinary Sciences<br>Utrecht University<br>Yalelaan 2, 3584 CM<br>Utrecht, The Netherlands |
| 1.5. Primary contact name and email address | Prof. Dr. M.H.M. Wauben<br> |
| 1.6. Date or time period of experiment | March 2018 – September 2019 |
| 1.7. Conclusions | LPP can be stained with PKH67 and detected upon fluorescence-based flow cytometry. The fluorescence and light scatter signals from stained LPP are in range to those of isolated and purified EV. Furthermore, when LPP were spiked-in presence of EV, the staining and detection of EV was affected.<br>The detection of an EV-marker CD9 was influenced in presence of LPP, leading to an underestimation of the number of CD9+ EV. |
| 1.8. Quality control measures | Procedural control for fluorescent stainings<br>Matched isotype PE control<br>Serial dilutions<br>Detergent treatment control |
| 2.1.1.1. (2.1.2.1., 2.1.3.1.) Sample description | EV isolated from 4T1 cell culture supernatant, Commercially available Lipoprotein Particles |
| 2.1.1.2. Biological sample source description | Cell culture supernatant from the murine mammary carcinoma cell line 4T1 (ATCC, Manassas, VA) was used to isolate EV.<br>Purified human lipoprotein particles: |

| | |
|---|---|
| | human chylomicrons (Catalog no. 7285-1000, Biovision Incorporated)<br>human very-low-density (Catalog no. 437647-5MG, Merck Millipore)<br>human low-density (Catalog no. LP2-2MG, Merck Millipore) |
| 2.1.1.3. Biological sample source organism description | Mouse and human |
| 2.1.2.2. Environmental sample location | |
| 2.3. Sample treatment description | For detergent treatment control samples were treated with 0.1% (v/v) triton X-100 (SERVA Electrophoresis GmbH, Heidelberg, Germany) final concentration for 30 seconds before reanalysis. |
| 2.4. Fluorescence reagent(s) description | PKH67 (Sigma-Aldrich)<br>Rat anti-mouse CD9PE Clone: KMC8, IgG2a, k (Lot No. 7268877, Becton Dickinson Biosciences)<br>Isotype Rat IgG2a, k PE (Lot. No. 8096525, Becton Dickinson Biosciences) |
| 3.1. Instrument manufacturer | Becton Dickinson |
| 3.2. Instrument model | BD Influx™ optimized instrument for detection of sub-micron sized particles as described previously. |
| 3.3. Instrument configuration and settings | BD Influx optimized to measure small particles. All configuration details can be found in van der Vlist, E. J., Nolte-'t Hoen, E. N., Stoorvogel, W., Arkesteijn, G. J., and Wauben, M. H. (2012) Fluorescent labeling of nano-sized vesicles released by cells and subsequent quantitative and qualitative analysis by high-resolution flow cytometry. Nat Protoc 7, 1311-1326.<br>Briefly, samples were measured at a constant flow rate for 30 seconds using a fluorescence threshold on the 488 nm laser. Threshold level was set to detect 10-20 events per second when measuring a buffer control. PE signals obtained upon excitation with the 561 nm laser were collected in the 585/42 band pass filter. |
| 4.1. List-mode data files | Are available upon request. |
| 4.2. Compensation description | No compensation was required due to instrument configuration. |
| 4.3. Data transformation details | No data transformation was applied. |
| 4.4.1. Gate description | Gates were applied based on the controls or the matched isotype control for the specific staining with CD9 antibody (shown in Supplementary Material & Methods). |

| | |
|---|---|
| 4.4.2. Gate statistics | The number of total events recorded in 30 seconds measurements are shown in the dot plots without any background correction.<br>GeoMean of PE of the total number or gated events are shown and respectively described. |
| 4.4.3. Gate boundaries | Images of the ungated and gated plots can be found in figures and supplementary data. |

**Supplementary Table 2. MIFlowCyt-EV framework.**

| | |
|---|---|
| 1.1 Preanalytical variables conforming to MISEV guidelines | Yes, all relevant data has been submitted to EV-TRACK for transparent reporting and centralizing knowledge in extracellular vesicle research (EV-TRACK ID: EV190078). |
| 1.2 Experimental design according to MIFlowCyt guidelines | Yes, MIFlowCyt checklist can be found as part of the supporting information of this manuscript. |
| 2.1 Sample staining details | Yes, described in Materials and Methods. |
| 2.2 Sample washing details | Yes, described in Materials and Methods. |
| 2.3 Sample dilution details | Yes, described in Materials and Methods. |
| 3.1 Buffer-only controls | Yes, relevant buffer controls were measure and are shown. |
| 3.2 Buffer with reagent controls | Yes, see Figure S2c and S2d. |
| 3.3 Unstained controls | N/A |
| 3.4 Isotype controls | Yes, see Figure S2d. |
| 3.5 Single-stained controls | N/A |
| 3.6 Procedural controls | Yes, see Figure S2c and d. |
| 3.7 Serial dilutions | Yes, serial dilutions were performed in previous characterization experiments to determine the ideal dilution used in this study. See Figure S2b. |
| 3.8 Detergent-treated controls | Yes, sensitivity to triton X-100 was determined in previous characterization experiments. See Figure S2 c and d. |
| 4.1 Trigger channel(s) and threshold(s) | Yes, all relevant details can be found in Materials and Methods. |
| 4.2 Flow rate / volumetric quantification | Yes, low flow rate was kept constant and is described in Materials and Methods. |
| 4.3 Fluorescence calibration | Yes |
| 4.4 Scatter calibration | N/A |
| 5.1 EV diameter/surface area/volume approximation | N/A |
| 5.2 EV refractive index approximation | N/A |
| 5.3 EV epitope number approximation | N/A |
| 6.1 Completion of MIFlowCyt checklist | Yes, see Table S1 |
| 6.2 Calibrated channel detection range | 100 FITC MESF |

| | |
|---|---|
| 6.3 EV number/concentration | Yes, see Figure 1. |
| 6.4 EV brightness | Yes, reported in ERF or MESF, see Figure 4d. |
| 7.1 Sharing of data to a public repository | Yes, all experimental details about the biological sample preparation can be found in EV-TRACK. All data files are available upon request |
